# UVA radiation could be a significant contributor to sunlight inactivation of SARS-CoV-2

**DOI:** 10.1101/2020.09.07.286666

**Authors:** Paolo Luzzatto-Fegiz, Fernando Temprano-Coleto, François J. Peaudecerf, Julien R. Landel, Yangying Zhu, Julie A. McMurry

## Abstract

Past experiments demonstrated SARS-CoV-2 inactivation by simulated sunlight; models have considered exclusively mechanisms involving UVB acting directly on RNA. However, UVA inactivation has been demonstrated for other enveloped RNA viruses, through indirect mechanisms involving the suspension medium. We propose a model combining UVB and UVA inactivation for SARS-CoV-2, which improves predictions by accounting for effects associated with the medium. UVA sensitivities deduced for SARS-CoV-2 are consistent with data for SARS-CoV-1 under UVA only. This analysis calls for experiments to separately assess effects of UVA and UVB in different media, and for including UVA in inactivation models.

**Lay summary:** Recent experiments have demonstrated that SARS-CoV-2 is inactivated by simulated sunlight; however, there are still many unknowns, including the mechanism of action and which part of the light spectrum is principally responsible. Our analysis indicates the need for targeted experiments that can separately assess the effects of UVA and UVB on SARS-CoV-2, and that sunlight inactivation models may need to be expanded to also include the effect of UVA. A first UVA-inclusive model is also proposed here. These findings have implications for how to improve the safety of the built environment, and for the seasonality of COVID-19.

## Background

Predicting the loss of infectivity of SARS-CoV-2 in various environmental conditions is important to mitigate the current COVID-19 pandemic. Laboratory studies found that although the virus persists at least for several hours in darkness and at room temperature [1], simulated sunlight induces rapid inactivation [2,3]. This is consistent with epidemiological studies on effects of UV [4], which is the wavelength band assumed to be responsible for inactivation [5].

Ratnesar-Shumate et al. [2] examined SARS-CoV-2 in simulated saliva and in complete growth medium (gMEM). The suspensions were applied to stainless steel surfaces and exposed to simulated sunlight. Inactivation increased with light intensity, and was faster in simulated saliva than in gMEM. Schuit et al. [3] aerosolized SARS-CoV-2 in the same two suspension media as in [2], varying humidity and sunlight intensity. They found weak dependence on humidity, but strong sensitivity to sunlight and medium. In both studies [2,3], the combination of sunlight and simulated saliva caused faster inactivation than was observed for sunlight and growth medium, although no significant difference between the two media was observed in darkness.

A model of SARS-CoV-2 sunlight inactivation would be highly valuable. Lytle & Sagripanti [6] modeled sunlight inactivation for viruses of interest to biodefense; noting that UVC radiation does not reach the Earth’s surface, they assumed an inactivation mechanism relying on UVB-induced damage to viral RNA (or DNA). Their model predicts inactivation rates from a given UVB irradiance spectrum and genome size. Sagripanti & Lytle [7] later calculated inactivation rates for SARS-CoV-2, for several worldwide locations at different times of the year. As will be shown later in Figure 1, the experimentally observed inactivation rates in [2,3] are significantly faster than predicted by the model in [6,7]. In addition, the observed dependence on the medium composition is not consistent with this medium-independent model.

**Figure 1.**
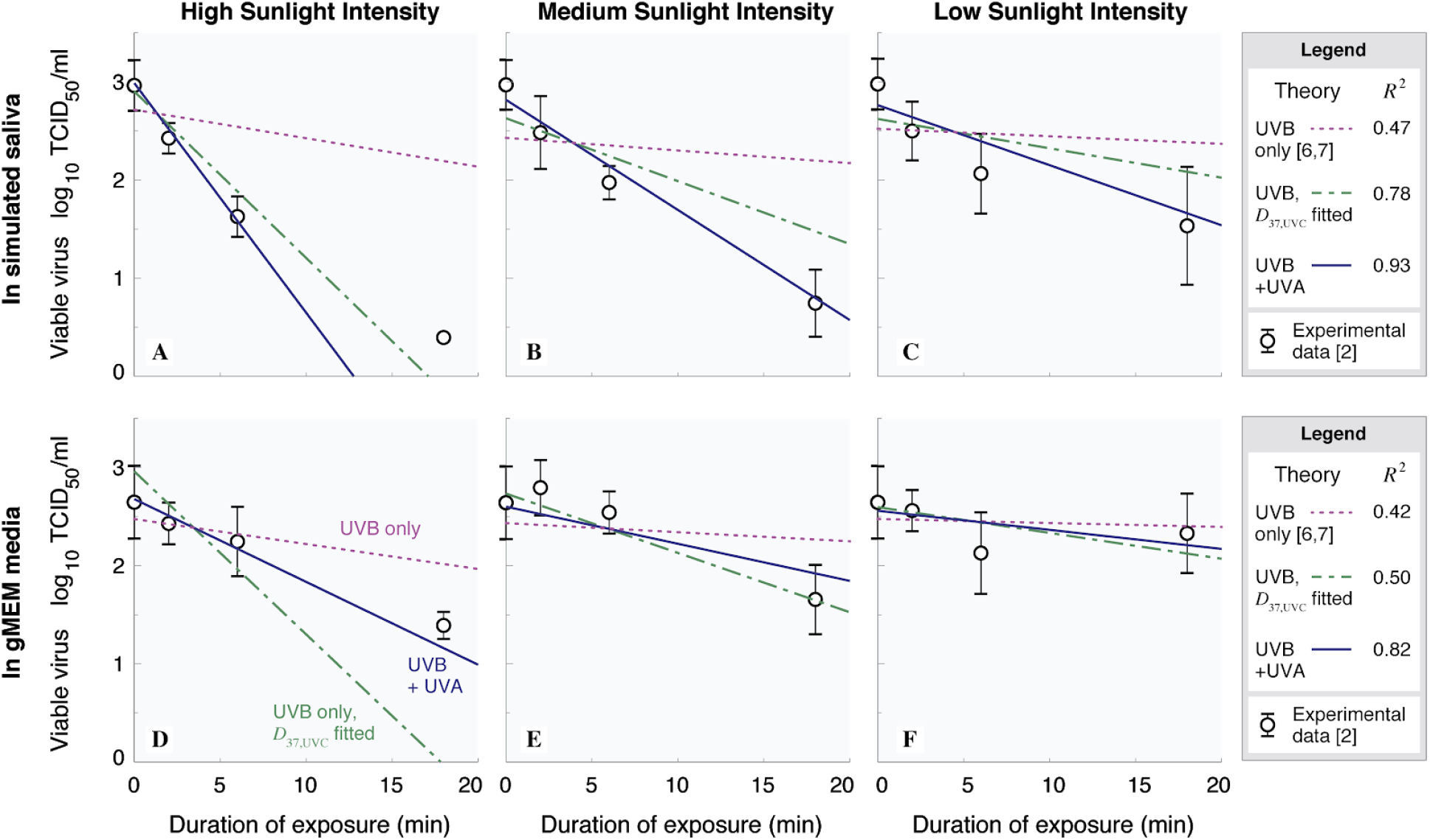
Comparison of theories with data for SARS-CoV-2 on stainless steel surfaces. (A, B, C): simulated saliva, (D, E, F): gMEM (complete growth medium). (A, D): high simulated sunlight, (B, E): medium simulated sunlight, (C, F): low simulated sunlight. Purple dotted line: UVB-only theory (Equation 3) with D_37,UVC_ = 3.0 J/m^2^ [6,7]. Green dot-dashed line: UVB-only theory with D_37,UVC_ from a fit to all data. Solid line: present model, combining UVB model (with D_37,UVC_ = 3.0 J/m^2^) and UVA model, with D_37,UVA_ from fits using all data for each medium. Symbols: data of Ratnesar-Shumate et al. [2]; for clarity, averages at each time are plotted, with error bars showing standard deviation.

Besides direct nucleic acid damage, other mechanisms of sunlight inactivation are possible, as reviewed by Nelson et al. [5] and illustrated in Supplementary Figure 1, also deposited in [8]. UV inactivation is first characterized as either endogenous or exogenous, that is, affecting the interior or the exterior of the virus. Second, inactivation can be either direct or indirect, that is, able to immediately damage the virus, or relying on an intermediate step. Indirect inactivation involves sensitizer molecules, generally provided by the medium, whose interaction with UV yields photo-produced reactive intermediates that can damage the virus [9].

It is generally expected that UVB acts on viruses through a direct, endogenous pathway [5], as has been modeled by [6,7]. Inactivation by UVA has been established for viruses in presence of sensitizers, through an exogenous, indirect pathway, for example by damaging the virus’s envelope and thereby disrupting attachment and entry [5,9]. Mechanisms that use other pathways (endogenous and indirect, or exogenous and direct) are deemed rare in viruses, since they contain few internal or surface sensitizers that absorb sunlight [5].

To the best of our knowledge, SARS-CoV-2 has not yet been studied under UVA-only; however, Darnell et al. [10] examined SARS-CoV-1 under exposure to either UVC or UVA over 15 minutes in Dulbecco’s modified Eagle’s medium (DMEM) with supplements. They found a clear effect of UVC, whereas inactivation due to UVA was of the order of measurement uncertainty. Darnell & Taylor [11] later found that UVA irradiation of SARS-CoV-1 in DMEM provided appreciable inactivation after 30 minutes; moreover, adding a photosensitizer inactivated the virus below the limit of detection within 15 minutes.

These existing results point towards a vulnerability of SARS-CoV-2 to UVA in the presence of sensitizers. Since the relative intensities of UVA and UVB vary greatly in sunlight, establishing UVA sensitivity of SARS-CoV-2 is necessary to obtain practically valuable predictions. In addition, since UVA from sunlight is abundant over a broader range of dates and times than UVB, vulnerability to UVA would imply a much greater sunlight inactivation potential than currently expected. This vulnerability could enable disinfection using inexpensive and efficient UVA light sources. This study tested the hypothesis that a model inclusive of UVA could improve agreement with SARS-CoV-2 experiments, with UVA sensitivities consistent with those for other similar viruses.

## Methods

Even without irradiation, the decay in infectious virion concentration over time is traditionally expressed as an exponential with a medium-dependent rate [12]. Here this rate is written as *k*_0_^medium^, where “medium” can be either simulated saliva or growth medium [2,3]. Therefore the infectious virion concentration *V* changes with time *t* following the expression

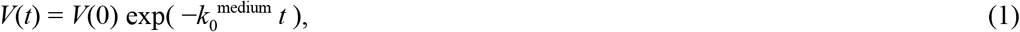

where *V*(0) is the initial concentration. The value of *k*_0_^medium^ is obtained from fit to data, and is negligible compared to sunlight effects [2,3].

To model sunlight inactivation in a general manner, one needs a hypothesis for the inactivation mechanism. As mentioned earlier, Lytle & Sagripanti [6] considered direct photochemical damage to viral RNA, which is maximal for UVC (wavelengths between 200-280 nm). UVC sensitivity was expressed as the exposure, at 254 nm, that produces one e-fold reduction in infectious virion concentration (i.e. to 37% of the initial value), and is written as *D*_37,UVC_ [6]. Based on genome size, for *Coronaviridae*, they estimated *D*_37,UVC_ between 2.5-3.9 J/m^2^, and *D* = 3.0 J/m^2^ for SARS-CoV-2 [7]; this value is used in the calculations presented here. Although no UVC reaches the Earth’s surface, longer wavelengths can still affect viral RNA, albeit with decreased sensitivity. To account for this, Lytle & Sagripanti [6] introduced a relative sensitivity, expressed as the ratio between sensitivity at a given wavelength *λ* and the UVC sensitivity at 254 nm [6] (see Supplementary Figure 1). Writing this relative sensitivity as *r*(*λ*), and expressing the spectral irradiance at a given wavelength as *E*_e,λ_(*λ*), one can evaluate an “equivalent UVC” irradiance (in W/m^2^) as

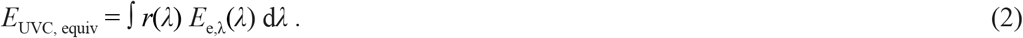

Since *r*(*λ*) drops to around 10^−4^ by a wavelength of 320 nm, this integral is performed only over the UVB spectrum (280 to 315 nm). In the calculations reported here, the irradiance spectra of [2,3] are used for *E*_e,λ_(*λ*), and the integral is performed numerically. For a dried thin layer on a surface, or for an aerosol particle, one might assume that the medium provides negligible light attenuation; inactivation by direct photochemical damage to RNA would therefore be independent of medium, and the infectious virion concentration would decay as

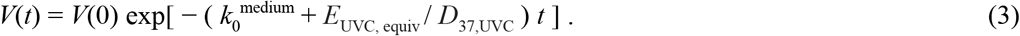

Here, an extension of this model is proposed, to account for indirect inactivation of SARS-CoV-2 due to UVA (315 to 400 nm). The integrated energy across the UVA range is considered. Although a specific wavelength may have relatively stronger impact, this a reasonable approach for sunlight, since the solar spectrum is relatively flat in this range (compared to UVB), such that the integrated energy is a proxy for the energy at any one wavelength, as discussed further below.

The UVA energy that produces an e-fold reduction in *V* is labeled *D*_37,UVA_^medium^, dependent on the medium, and the irradiance integrated across the UVA range is *E*_UVA_. Therefore the hypothesis being tested here is that, under sunlight, the infectious virion concentration is approximated by

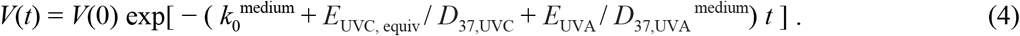

Note that this modeling approach is also consistent with established theories of photochemical kinetics [13], which are widely used (including e.g. for photochromic molecules [14]).

Since *D*_37,UVA_^medium^ is medium-dependent, it is found by fitting experimental data. For experiments that report infectious virion concentrations, nonlinear model fitting in MATLAB is used, on a logarithmic scale. For experiments that only report an overall inactivation rate at each irradiation condition, *D*_37,UVA_^medium^ is calculated at each condition and averaged over experiments with the same medium.

## Results

The infectious virion concentration data of Ratnesar-Shumate et al. [2] is shown in Figure 1, in units of tissue culture infectious dose/ml (TCID_50_/ml). For clarity, instead of plotting all data points, a circle shows the average at each time, and an error bar shows the standard deviation.

The model based only on UVB inactivation [6,7], embodied by Equation (3) with *D*_37,UVC_ = 3.0 J/m^2^, is shown by the purple dotted line, and had an R^2^ (computed across all light intensities) of 0.47 for simulated saliva and 0.42 for gMEM. To test whether this could be improved by a different value of *D*_37,UVC_, a fit was sought across all data (since endogenous inactivation is expected to be medium-independent), and is shown by the green dot-dashed line in Figure 1. The fit yields *D*_37,UVC_ = 0.53 ± 0.11 J/m^2^ (where the uncertainty corresponds to the 95% confidence interval), with updated R^2^ values of 0.78 and 0.50. This *D*_37,UVC_ is at least five times smaller than expected for any coronavirus (2.5 - 3.9 J/m^2^), or for any virus considered by [6]. This suggests that SARS-CoV-2 is more sensitive to sunlight than expected by UVB-induced, endogenous inactivation alone.

The model inclusive of UVA, described by equation (4), is plotted by the solid blue line in Figure 1. The value of *D*_37,UVA_ is found by fitting separately for each medium, consistently with the exogenous, indirect inactivation hypothesis. This yields *D*_37,UVA_ = 10 ± 1.8 kJ/m^2^ for simulated saliva and *D* = 30 ± 14 kJ/m^2^ for gMEM. The updated values of R^2^ (computed across all light intensities) are 0.93 and 0.82 for simulated saliva and gMEM respectively. Applying this theory also to the aerosol data of Schuit et al. [3] gives *D*_37,UVA_ = 34 ± 11 kJ/m^2^ in simulated saliva and *D* = 88 ± 50 kJ/m^2^ for gMEM; these appreciable uncertainties seem to reflect the relatively small number of replicates available (for comparison, the influenza experiments of [12] yielded a smaller uncertainty, as shown below).

To assess whether these *D*_37,UVA_ values are plausible, data for UVA-only inactivation of SARS-CoV-1 was also examined. For DMEM with supplements, [10] implies *D*_37,UVA_ = 14 ± 10 kJ/m^2^ (data was collected only over 15 minutes, making it difficult to reduce uncertainty). For DMEM without supplements, [11] implies *D*_37,UVA_ = 9.2 ± 4.2 kJ/m^2^. To provide additional context, *D* was calculated also for influenza, which like SARS-CoV-2 is an enveloped, RNA virus. Data of [12] was used for influenza, which examined a gMEM-based aerosol under simulated sunlight. Using *D*_37,UVC_ = 7.3 J/m^2^ from [6], one deduces *D*_37,UVA_ = 12 ± 2.0 kJ/m^2^. These results are summarized in Table 1.

**Table 1.**
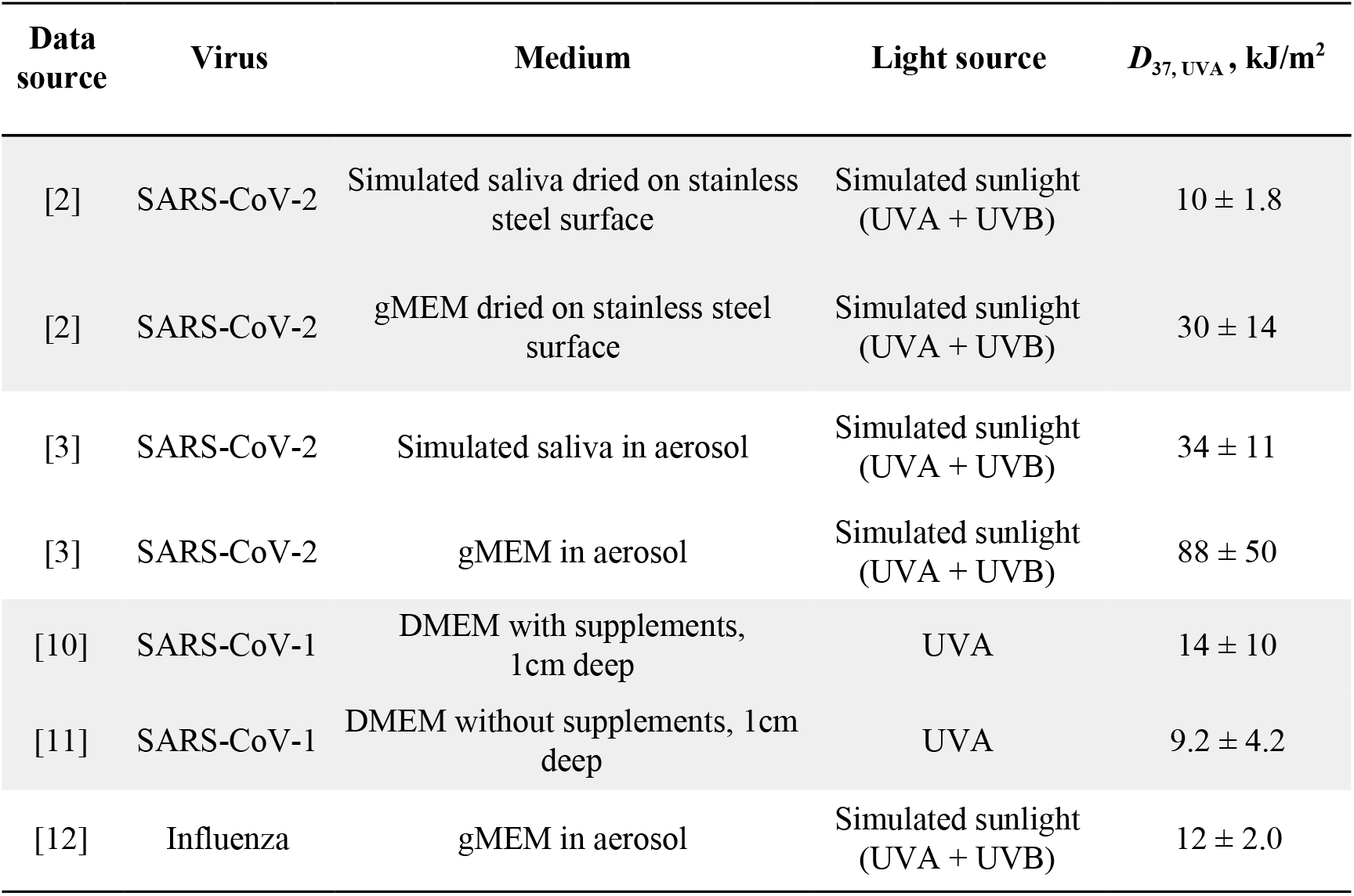
Summary of UVA sensitivities deduced from published data. Sensitivity to UVA for a given medium and virus. The parameter D_37,UVA_, computed by fitting the proposed model (Equation 4) with the data in Figure 1, indicates the amount of energy required to achieve one e-fold reduction in virion concentration. Smaller D_37,UVA_ indicates that less energy is required for inactivation, and therefore that the virus has greater sensitivity. DMEM: Dulbecco’s modified Eagle’s medium; gMEM: complete growth medium.

| Data source | Virus | Medium | Light source | $D_{37, UVA}$, kJ/m<sup>2</sup> |
| --- | --- | --- | --- | --- |
| [2] | SARS-CoV-2 | Simulated saliva dried on stainless steel surface | Simulated sunlight (UVA + UVB) | $10 \pm 1.8$ |
| [2] | SARS-CoV-2 | gMEM dried on stainless steel surface | Simulated sunlight (UVA + UVB) | $30 \pm 14$ |
| [3] | SARS-CoV-2 | Simulated saliva in aerosol | Simulated sunlight (UVA + UVB) | $34 \pm 11$ |
| [3] | SARS-CoV-2 | gMEM in aerosol | Simulated sunlight (UVA + UVB) | $88 \pm 50$ |
| [10] | SARS-CoV-1 | DMEM with supplements, 1cm deep | UVA | $14 \pm 10$ |
| [11] | SARS-CoV-1 | DMEM without supplements, 1cm deep | UVA | $9.2 \pm 4.2$ |
| [12] | Influenza | gMEM in aerosol | Simulated sunlight (UVA + UVB) | $12 \pm 2.0$ |

## Discussion

These calculations showed that a model based only on a UVB-driven, endogenous direct mechanism is not sufficient to explain SARS-CoV-2’s rapid decay and medium sensitivity under simulated sunlight. To account for these discrepancies, a model inclusive also of exogenous, indirect inactivation due to UVA was proposed and tested (Equation 4). This model improved agreement with data across all conditions. Moreover, the UVA sensitivities deduced for SARS-CoV-2 were comparable to those found for SARS-CoV-1 exposed only to UVA, as well as for influenza.

Greater UVA sensitivity was found on stainless steel surfaces than in aerosols. This could have several possible explanations; for example, surface reflectivity could effectively increase irradiation by up to 40% [15]. However, since the uncertainty from aerosol experiments is larger than for surfaces, more data is needed to confirm significant differences.

These results point to the need for experiments to separately test effects of UVA, UVB, and of medium composition. Key sensitizers may vary between simulated and real saliva, as well as across saliva from different individuals, leading to different inactivation rates. Sensitizers may respond to narrower wavelength ranges than the full UVA band considered in our model. If UVA sensitivity is confirmed, sunlight could mitigate outdoor transmission over a broader range of latitudes and daytimes than previously expected, since UVA is less strongly absorbed by atmospheric ozone than UVB. For example, at midlatitudes, UVA changes approximately by a factor of two between summer and winter, whereas UVB varies by a factor of four [5]. Furthermore, inexpensive and energy-efficient UVA sources might be used to augment air filtration systems at relatively low risk for human health, especially in high-risk settings such as hospitals and public transportation.

## Meetings where the information has previously been presented

This work appeared as a poster presentation in the meeting “Climatological, Meteorological, and Environmental factors in the COVID-19 pandemic”, 4-6 August 2020. https://public.wmo.int/en/events/meetings/covid-19-symposium

